# Impacts of Anthropogenic, Environmental, and Genetic Factors on Peanut Seed Protein Content and Composition

**DOI:** 10.64898/2026.10.17.759842

**Authors:** Hrishikesh P. Ingole, Zachary T. Jones, Sachin Rustgi

**Affiliations:** Department of Plant and Environmental Sciences, Clemson University Pee Dee Research and Education Center, Florence, SC, USA

**Author notes:** Authors contributed equally to this research (Hrishikesh P. Ingole, Zachary T. Jones).

**Keywords:** Peanut, domestication, breeding, seed storage proteins, allergenicity, homoeologous exchange

## Abstract

Peanut contributes approximately $4 billion annually to the U.S. economy, with 13 peanut-producing states generating over 3 mmt, 75–80% of which is consumed domestically. Peanut seeds contain 22–36% protein and 40–56% oil, enriched in unsaturated fatty acids, making them a highly nutritious food source. However, specific seed storage proteins can trigger allergic reactions ranging from mild symptoms to severe anaphylaxis, underscoring the need to better understand factors influencing seed protein composition. Anthropogenic processes such as domestication and modern breeding, environmental stresses including excessive heat during flowering and pod development, and genetic mechanisms such as homoeologous exchanges (HEs) collectively shape seed protein content and composition. Domestication improved harvestability, while breeding enhanced yield and nutritional quality. Current improvement programs increasingly emphasize heat resilience in response to a ~1.5 °C rise in global temperatures, with further warming anticipated. Yet limited attention has been given to how these factors affect protein composition, allergenic potential, trait stability under future climates, and genotypic uniformity linked to HEs. Here, we review the impacts of domestication, breeding, environmental stress, and genomic restructuring on peanut seed protein content and composition, highlighting key research gaps to ensure nutritional quality, safety, and stability in a changing climate.

**Main conclusion:** Our research ruled out domestication and modern breeding as drivers of increased peanut allergen content and allergy prevalence, identifying homoeologous recombination and heat stress as factors affecting seed protein composition.

## Introduction

Peanut or groundnut (*Arachis hypogaea* L.) evolved as a tetraploid species (AABB, 2*n* = 4× = 40) from a single hybridization event between two wild diploid progenitors, *Arachis duranensis* (A-subgenome donor) and *Arachis ipaensis* (B-subgenome donor), 5,000-10,000 years ago at an unknown site in the southern Bolivia–northwestern Argentina region (Kochert et al., 1996; Stalker, 2017). Since its first cultivation in northern Peru 8,500 years ago, peanut has become a major global crop, with 51.4 mmt produced worldwide (Hammons, 1973; Dillehay et al., 2007; USDA FAS, 2024), and the third-largest source of plant-based protein (Savage and Keenan, 1994; Cui et al., 2023).

Nearly one-third of the world’s population suffers from protein malnutrition and diseases associated with poor protein quality (Roth, 2013). Therefore, selecting for high protein content has remained an important breeding objective and a general preference. However, a critical evaluation of seed protein composition is necessary, as approximately 2% of the Western population is sensitive to peanuts, with 17 seed proteins implicated in causing allergic reactions (Shah et al., 2019; Yu and Eghbali, 2025). Specifically, storage proteins such as arachin (legumin-like, ~12S globulin), conarachin (vicilin-like, ~7S globulin), and the smaller 2S albumin—together representing 37-45% of total seed proteins—are the primary etiological agents (Dreskin et al., 2021; Rustgi et al., 2022). Among these, Ara h1, Ara h2, Ara h3, and Ara h6 are the most prevalent in eliciting allergic responses (Shah et al., 2019; Alam and Rustgi, 2025).

Seed storage proteins (SSPs) in peanut accumulate predominantly in the embryo, specifically the cotyledons, during seed filling and maturation (Kang et al., 2007). The expression of SSP-encoding genes is cotyledon-specific and is regulated by a complex network of cis-regulatory elements and trans-acting factors (Vicente-Carbajosa and Carbonero, 2005). Recent research has further shown that the expression and protein abundance of major peanut allergens, measured by IgE binding, increase progressively throughout seed maturation (Cohen et al., 2024). This regulatory network comprises proximal and distal promoter elements, transcription factors and repressors, components of the transcriptional machinery, epigenetic regulators, and proteins involved in SSP processing and transport (Yang et al., 2023). The regulatory mechanisms controlling SSP accumulation have been extensively characterized in *Arabidopsis* (Brassicaceae) and soybean (Fabaceae) (reviewed in Yang et al., 2023). In contrast, comparable studies in peanut (Fabaceae) remain limited, and much of the current understanding has been inferred from soybean. Nevertheless, the *Ara h1* promoter was recently cloned and shown to contain seed-specific RY motifs that drive seed- and cotyledon-specific expression in heterologous *Arabidopsis* systems (Yuan et al., 2022). More recently, our group used genome-wide association mapping to identify cis-and trans-regulatory loci associated with the accumulation of the major peanut allergens (Ara h1, Ara h2, Ara h3, and Ara h6) (Alam et al., 2026). Ongoing studies are further elucidating the regulatory mechanisms underlying allergen accumulation to improve our understanding of peanut allergome composition and its regulation during seed development.

Over the years, peanut domestication and breeding efforts have primarily focused on improving harvestability and increasing yield by enhancing resistance to abiotic and biotic stressors while also developing desirable seed oil profiles (Holbrook et al., 2016) and increased total seed protein content (Li et al., 2025). However, relatively little attention has been given to how these manipulations have affected the abundance and composition of immunogenic proteins. This raises an important question: have humans inadvertently increased the allergenic potential of peanuts through domestication and modern breeding, particularly in light of the rising prevalence of peanut allergy (Warren et al., 2021)?

Addressing this question requires comprehensive studies of seed protein content and composition, together with a better understanding of the complex genetic mechanisms regulating these traits. Cultivated peanut is a relatively young allotetraploid that underwent a severe genetic bottleneck during its evolution. Despite its limited genetic diversity, it generates novel variation through tetrasomic recombination, or homoeologous chromosome exchanges (Lamon et al., 2025a, b). This process creates genetic heterogeneity even within otherwise genetically uniform lines, posing a unique challenge when evaluating biomolecular traits such as seed protein composition from individual seeds. In contrast, agronomic and physiological traits are typically measured at the plot level, where phenotypic variation among individual plants is averaged, thereby reducing the impact of within-line heterogeneity.

Furthermore, understanding how environmental factors influence seed protein content and composition is essential for assessing the stability of these traits under changing climatic conditions. To address these knowledge gaps, this study investigated the effects of domestication and breeding, environmental stressors (particularly heat stress), and genomic instability resulting from homoeologous chromosome exchanges on allergen content in peanut seeds.

### Impact of human intervention on peanut seed protein content and composition

The domestication of peanuts can be traced through a complex history that integrates genetic evolution with human influences. Domestication is often characterized by polyploidization events and mutations that lead to the rapid accumulation of favorable agricultural traits, such as those observed in peanut (Zhang et al., 2017; Leal-Bertioli et al., 2021). As mentioned, peanut originated through a single hybridization event between closely related A- and B-subgenome progenitors (with over 98% DNA identity between corresponding genes and retention of approximately 90.68% of orthologous gene families), resulting in a massive genetic bottleneck that constrained the array of possible allelic combinations associated with beneficial agronomical traits, such as resistance to pests and pathogens (Chu et al., 2021, Bertioli et al., 2019, 2020). This understanding has prompted researchers to mine the diploid ancestors of peanut for more favorable alleles, with the goal of increasing genetic diversity and incorporating desirable alleles into the tetraploid genome (Stalker, 2017). This effort is most notably demonstrated by the introgression of *Arachis cardenasii* and *A. diogoi* into the tetraploid peanut to enhance disease resistance (Bertioli et al., 2021; Hancock et al., 2019).

Inspired by the earlier findings, the authors evaluated *A. hypogaea* relatives for seed protein content and composition in the hope of identifying peanuts with reduced immunogen content. Altogether, six A subgenome and three B subgenome peanut species, a synthetic tetraploid (IpaDur1, PI 707937), and a wild tetraploid *Arachis monticola*, were evaluated for total seed protein content and composition using the Thermo Scientific Pierce BCA (bicinchoninic acid) kit and sodium dodecyl sulfate–polyacrylamide gel electrophoresis (SDS PAGE), as outlined in Alam et al. (2026). A larger representation of the A subgenome lines was reflective of the proportion of lines with different subgenomes in the U.S. peanut germplasm collection (Bertioli et al., 2021). Electrophoretic analysis revealed that none, other than a single *A. diogoi* accession (GKP 10602, ICG 4983, PI 276235), displayed reduced accumulation of Ara h1 and Ara h3 relative to the tetraploid *A. hypogaea* cultivar ‘Gregory,’ (Fig. **1a**). which was independently and parallelly confirmed by Singh et al. (2018). Comparison of the allergen accumulation profiles of the A and B subgenome progenitors and their synthetic hybrid IpaDur1 revealed gel patterns that resembled with each other’s and *A. monticola*’s, with no signs of an increase in allergen content. The reduced accumulation of allergenic proteins in *A. diogoi* led to the screening of 121 introgression lines of ‘Gregory’ × *A. diogoi* population (84 ILs from Hancock et al., 2019 and additional lines received from N.C. State University) and a germplasm line PI 598133 (ICGV-SM 86715) with *A. diogoi* introgression(s) (Moss et al., 1998). Various biochemical (densitometric analysis of SDS-polyacrylamide gels, HPLC, LC-MS) and immunological assays (ELISA, Western blot with allergen-specific antibodies and IgE derived from the sera of allergic individuals) revealed introgression lines with variable levels of protein accumulation, including lines with reductions well exceeding those of *A. diogoi* (Fig. **S1**). Overall, these findings suggest that the allergenicity of tetraploid peanut did not increase due to domestication or the incorporation of genetic material from any of the wild relatives and progenitors tested here. Instead, in the case of a specific *A. diogoi* accession (GKP 10602), allergenicity may actually be reduced.

**Figure 1.**
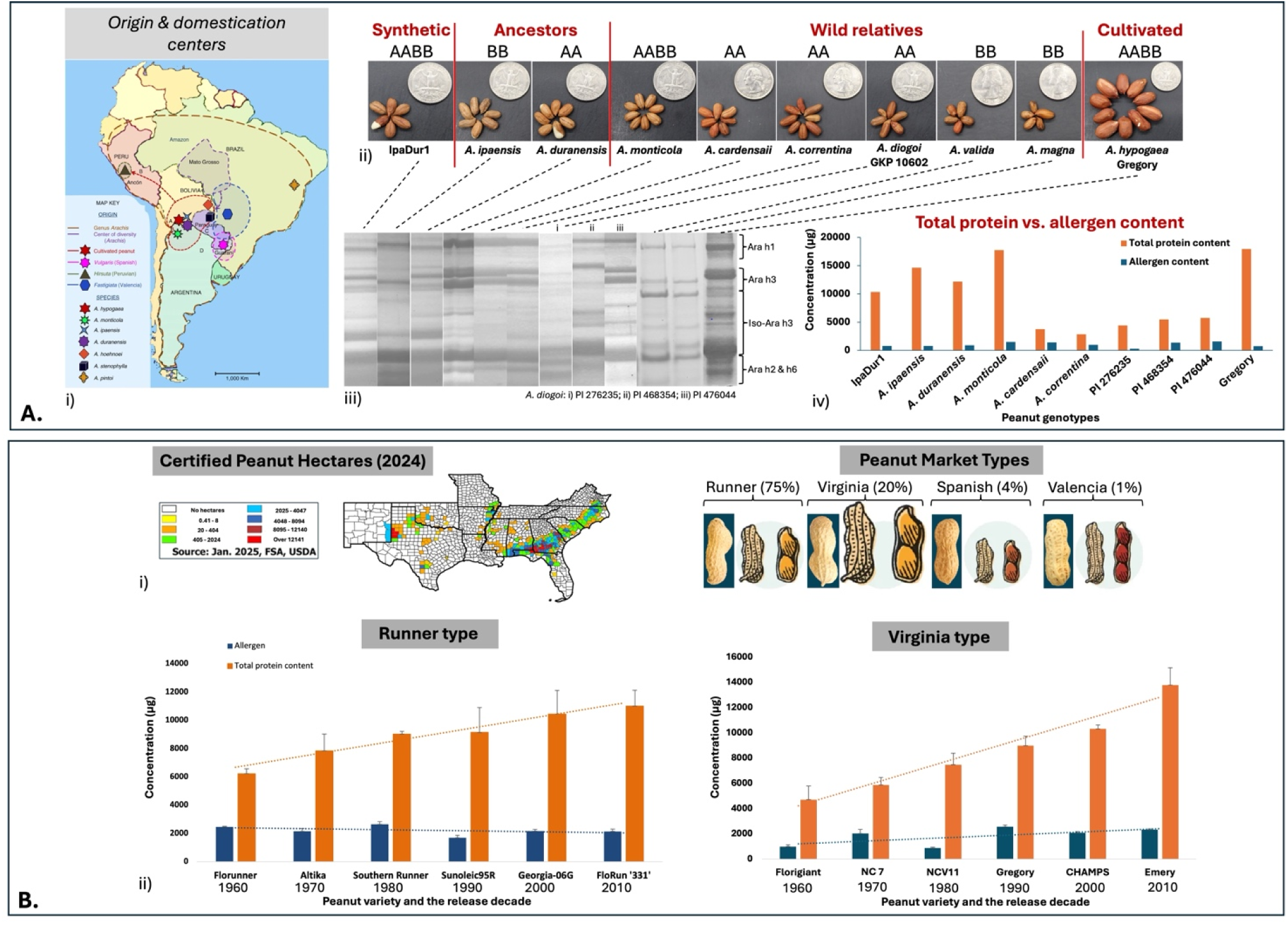
Evaluation of the impact of domestication (polyploidization) and breeding on seed protein content and composition. Panel A. Evaluation of diploid peanut progenitors, wild relatives, and a synthetic line (IpaDur1) for total seed protein content and composition using the Pierce BCA method and densitometric analysis of SDS polyacrylamide gels. i) Illustration of the peanut center of origin and domestication (adapted from 66 Zhuang et al., 2019). ii) Seed images of diploid peanut species (progenitors and wild relatives) and synthetic, wild, and cultivated tetraploid peanuts showing morphological differences with US quarter dollar for scale.. iii) SDS polyacrylamide gel profiles of seed storage proteins from wild (diploid and tetraploid), synthetic, and cultivated peanuts. iv) Quantification of total protein determined using the Pierce BCA assay and Ara h protein content determined through densitometric analysis of SDS polyacrylamide gels (relative quantification was performed using known quantities from a bovine serum albumin standard curve). Panel B. Examination of the relationship between total protein content and major allergen content in representative peanut varieties from two market classes (Runner and Virginia), released between 1960 and 2010, using the Pierce BCA method and densitometric analysis of SDS polyacrylamide gels. i) Illustration of peanut-growing areas in the southeastern United States (left) and the market share of different peanut classes within the U.S. ii) Total protein and allergen content of Runner and Virginia market classes

Modern breeding programs in peanuts have primarily focused on enhancing peanut yields by identifying and incorporating quantitative trait loci (QTLs) that influence yield-related traits. To the best of the authors’ knowledge, there are no studies that have demonstrated the long-term impact of peanut breeding programs on the allergen content. Thus, a panel of representative varieties from the Runner (75% of U.S. market) and Virginia (20% of U.S. market) peanut market classes for each decade from 1960 to 2020 was screened for total protein content using the Pierce BCA assay and densitometric analysis of polyacrylamide gels to determine allergenic protein content (Alam et al., 2026). The resulting allergenic protein values were compared with the total quantification of peanut protein, normalized to the seed weight. Interestingly, a comparison of the representative varieties in both the Virginia and Runner market classes suggested that total protein content increased over the years, while allergenic protein content did not follow the trend and remained consistent. Indicating that the U.S. breeding program strategies have not led to an increase in allergenic protein content (Fig. **1b**).

### Influence of Reproductive Heat Stress on Allergenic Protein Content in Peanuts

Heat stress in peanuts can have detrimental effects on yield, caused in part by a decrease in pollen viability, seed set, and biomass (Prasad et al., 1999; Ingole et al., 2025). Recent studies by Zoong Lwe et al. (2020) and Spivey et al. (2023) suggest that heat stress can alter leaf and anther lipid composition by remodeling cellular membrane lipids and fatty acid saturation levels, thereby preventing membrane hyperfluidization and the collapse of the lipid bilayer into a single layer. Heat stress during flowering can lead to changes in the amino acid profile of peanut seeds, resulting in reduced protein quality (Singh et al., 2016). Additional evidence in the oilseed crop *Camelina sativa* L. Crantz suggests that the timing of heat stress can have various effects on seed size and composition, dependent upon the plant’s reproductive stage at the time of exposure (Smith and Lu, 2024). Furthermore, Chakraborty et al. (2018) studied the impact of heat stress on heat-susceptible and heat-tolerant peanut genotypes and reported that the tolerant genotypes exhibited early induction of low–molecular weight heat shock proteins in leaf tissues relative to the susceptible genotypes. Likewise, Krishnan et al. (2020) reported an elevated accumulation of heat shock proteins during seed filling under heat stress in soybean, and that heat stress triggered a reduced accumulation of heat-stress–responsive proteins in seeds of susceptible genotypes, with minimal impact observed in tolerant genotypes. These findings led the authors to investigate whether the allergenic protein content of peanut will be impacted by potential increases in global temperatures.

A set of 12 peanut lines was selected for this experiment, which included six varieties representing decades 1960-2020—three Virginia and three Runner types, two recombinant inbred lines (RILs) with reduced allergen content from a ‘Georgia Green’ × N C 25 (PI 371521) population (N.C. State University), along with two varieties with contrasting levels of heat tolerance, ICGS 76 (heat-tolerant) and Tamrun OL 02 (heat-susceptible) (Spivey et al., 2023). Two additional lines, PI 356004 and PI 493581, were included due to their low and high pollen viability under heat stress, respectively (Ingole et al., 2025). The plants were subjected to heat stress at 38/26 °C (day/night) for 14 days in a growth cabinet, commencing at the onset of peg formation, as described by Spivey et al. (2023). Each line was sown in six replicates. Three replicates were exposed to heat stress, while the remaining three were maintained under optimal growth conditions. After heat exposure, plants were returned to normal temperature until pod harvest. Seeds from both treatments were analyzed for protein profiles using SDS-PAGE, with a focus on the four major peanut allergens—Ara h1, Ara h2, Ara h3, and Ara h6.

The seed protein content of peanut varieties exposed to heat stress was compared with that of plants grown under optimal conditions. The results revealed that protein content in ‘NC 7’ and ‘Florunner’ significantly increased under heat stress. The other varieties, except for ‘CHAMPS,’ showed a decrease in protein content under heat stress; however, the change was not statistically significant (Fig. **2**). In case of the two RILs 460 and 571 no significant differences were observed however 571 showed increase while 460 tend to decrease the protein content. The previously identified heat-tolerant genotype ‘ICGS76’ and susceptible ‘Tamrun OL 02’ showed no significant difference. In contrast, another heat-susceptible genotype, PI 356004—identified based on decreased pollen viability—showed a significant reduction in protein content under heat stress. Conversely, the high–pollen-viability line PI 493581, which is considered heat tolerant, exhibited a slight increase in protein content under heat stress (Fig. **S2**).

**Figure 2.**
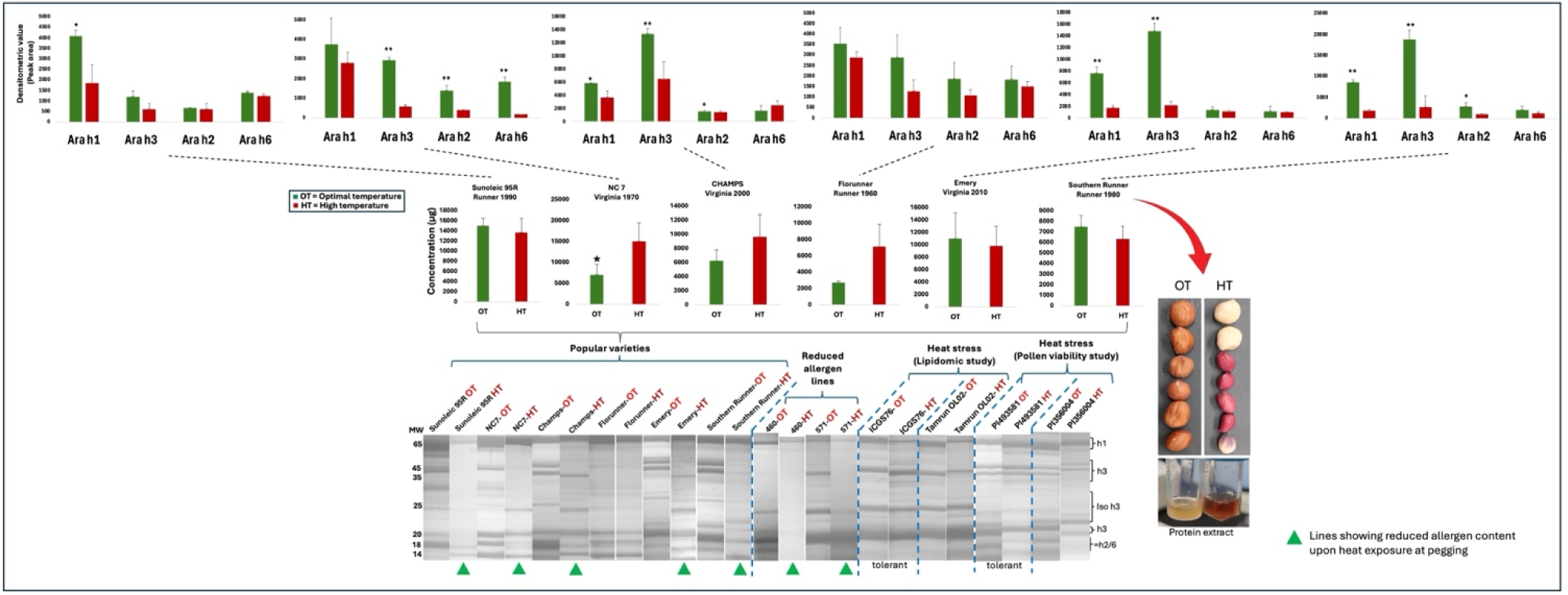
The impact of pegging stage heat stress on peanut seed storage protein composition. Determination of the effect of heat stress applied at the time of pegging in selected peanut varieties and germplasm lines, including lines with reduced allergen content and heat-tolerant lines. The top graphs show the levels of each immunogen (Ara h1, Ara h2, Ara h3, and Ara h6) in each line grown under optimal temperature conditions and subjected to heat stress at pegging. The middle graphs illustrate total seed protein content from extracts of optimally grown and heat-treated peanut plants of the corresponding varieties (bottom right highlights an observed seed coat color change in heat-treated Southern Runner seeds). The bottom panel presents SDS-PAGE images of selected popular varieties, reduced-allergen recombinant inbred lines from the NC 25 × ‘Georgia Green’ mapping population, and pre-identified heat-susceptible and heat-tolerant lines. Lanes marked with green triangles indicate a reduction in one or more Ara h proteins. * = p-value < 0.05; ** = p-value < 0.01.

Allergen content generally decreased under heat stress in the peanut varieties studied, except for ‘Florunner,’ which showed no significant change in the content of all four major allergens. For Ara h1, all varieties except ‘Florunner’ and ‘NC 7’ exhibited significant reductions; for Ara h3, ‘Southern Runner’, ‘CHAMPS’, ‘Emery,’ and ‘NC 7 showed notable decreases. For Ara h2, ‘Southern Runner’ and ‘CHAMPS,’ and ‘NC 7’ displayed significant reductions, whereas for Ara h6, a significant decline was observed only in ‘NC 7’ (Fig. **2**).

Among the two RILs, line 460 exhibited significant decreases in all allergens except Ara h1, while non-significant differences in the accumulation of allergenic proteins under heat stress and optimal growth conditions were observed for 571. The heat-susceptible, ‘Tamrun OL 02,’ showed minimal change, with Ara h1 being significantly reduced and Ara h3 significantly increased; in contrast, the tolerant genotype, ‘ICGS 76,’ demonstrated significant reductions in all four major allergens. The preidentified line with decreased pollen viability, PI 356004, displayed decreased total protein under heat stress, with Ara h1 and Ara h2 notably higher, whereas PI 493581, with high pollen viability, showed reduced allergen content under heat stress, significant only for Ara h3 and Ara h6 (Fig. **S2**).

Findings from this experiment indicate that heat stress can alter the protein composition of peanut seeds, and that other abiotic and biotic stressors may have similar effects. Although research on stress-induced changes in peanut and soybean remains limited, environmental stress responses have been extensively characterized in several legumes, including chickpea, common bean, faba bean, lentil, lupin, mung bean, and pea. Across these species, studies consistently show that primary metabolites—including seed and leaf proteins—as well as secondary metabolites can serve as useful biomarkers of heat and drought stress.

In faba bean, Abid et al. (2024) demonstrated that drought stress disrupts major physiological processes and triggers proline accumulation and increased antioxidant enzyme activity, including CAT, SOD, APX, and GPOX. Their proteomic survey identified 2,000 proteins, 81 of which were differentially expressed. These proteins were enriched in pathways associated with photosynthesis, ROS detoxification, amino acid and secondary metabolite biosynthesis, molecular chaperones, signal transduction, and energy metabolism, highlighting the complexity of the stress-responsive network. Similarly, Li et al. (2018) reported the differential expression of 50 proteins in drought-stressed faba bean seedlings, including the upregulation of chitinase and the downregulation of a 50S ribosomal protein. Using gel-based profiling, Abdellatif et al. (2012) further demonstrated that drought-tolerant and drought-susceptible faba bean varieties exhibit distinct seed-protein banding patterns.

Likewise, proteomic studies in cotton revealed that heat-tolerant genotypes maintain higher levels of proteins involved in photosynthesis, oxidoreductase activity, translation, transport, and carbohydrate and protein metabolism during high-temperature stress, whereas susceptible genotypes show widespread downregulation of these proteins (Perveen et al., 2025). In peanut, Katam et al. (2007) reported transcriptional and translational changes in response to drought stress. Using two-dimensional electrophoresis, mass spectrometry, and differential-display RT-PCR, they demonstrated that tolerant genotypes maintained relatively stable expression of methionine-rich seed-storage proteins, arachin, and conglutin, whereas drought-susceptible genotypes showed marked reductions in their accumulation.

Changes in amino acid composition under heat, drought, or salinity stress often result from protein degradation, shifts in protein synthesis, and the accumulation of free amino acids that function as osmolytes or ROS scavengers. For instance, under salinity stress, lysine levels typically decrease, whereas glycine, tyrosine, alanine, and isoleucine levels increase (Toderich et al., 2020). High-lysine barley mutants illustrate these changes, exhibiting reduced levels of hordein proteins and increased albumin and globulin fractions, particularly four lysine-rich, salt-soluble proteins: β-amylase, protein Z, and two chymotrypsin inhibitors (Munck, 1992). In hexaploid common wheat—particularly in diploid wheat species representing the A subgenome—and in maize, a correlation has been identified between grain lysine content and lysine-rich translation elongation factor 1-alpha (EF-1α), as well as other proteins associated with EF-1α. In wheat, one such protein was identified as fructose-1,6-bisphosphate aldolase. Immunological detection of these proteins was proposed as a potential marker of feed quality (Singh et al., 2000).

In addition to the abiotic factors discussed above, biotic factors such as viral infection can also influence seed-protein composition. For example, *Arabidopsis* infected with Tomato mosaic virus exhibited increased accumulation of vicilin-like proteins (Chen et al., 2013). More recently, the effects of soybean mosaic virus (SMV) infection on immunologically active proteins and on the seed accumulation of glycinin and β-conglycinin—both of which influence end-use performance—were investigated in conventional and transgenic soybean varieties. Infected seeds from the varieties examined showed increased levels of the highly immunoreactive Kunitz soybean trypsin inhibitor (KTI) and lectins, along with decreased flavonoid content, compared with healthy seeds. In addition, infected seeds exhibited reduced glycinin content and a lower glycinin-to-β-conglycinin ratio, although some varieties maintained their glycinin and β-conglycinin levels.

This pattern resembles the effects of heat stress on soybean seed-protein composition and drought stress on peanut seed composition. In both cases, tolerant varieties maintain relatively stable seed-protein profiles under stress, whereas susceptible varieties show reduced protein accumulation (Krishnan et al., 2020; Katam et al., 2007). These findings suggest that protein profiles could serve as biomarkers of resistance to viral infection (Molodchenkova et al., 2025).

Ongoing work is examining whether Tomato spotted wilt virus (TSWV) induces similar changes in peanut. In initial screenings of healthy and TSWV-infected seeds from ‘Georgia 16 HO,’ the authors observed no significant differences in protein or allergen content (Fig. **S3**). However, Xue et al. (2026) detected a sixfold increase in *Ara h13* (*DEF3*) expression 24 h after inoculation with *Ralstonia solanacearum* strain 180731-1, the causal pathogen of bacterial wilt, whereas no significant change in *Ara h12* (*DEF1*) expression was observed. These findings suggest that not all seed-storage proteins are affected equally by pathogenic infection.

In peanut, responses to infection and environmental stress have also been associated with seed-coat-mediated physical and biochemical barriers, particularly tannins—including proanthocyanidins—and other phenolic compounds. These compounds have been linked to differential susceptibility to aflatoxin contamination and to broader changes in nutritional quality (Puppala et al., 2023; Mendu et al., 2022). Further research is needed to fully characterize how pests and pathogens influence seed-protein content and composition and to elucidate the mechanisms underlying these effects.

### Protein biotypes and homoeologous recombination in peanut

Analysis of seed storage protein profiles across a diverse set of peanut germplasm—including popular varieties released over different decades, breeding lines and cultivars carrying heat or drought tolerance, resistance to aflatoxin accumulation, and accessions from the U.S. peanut mini-core collection—revealed that a large proportion of lines exhibited multiple protein biotypes when analyzed by SDS–PAGE (11.5% of lines in the U.S. peanut mini-core and 52.9% tested peanut varieties/breeding lines) (Fig. **S4**; Table **S1**). The possibility that this variation resulted from contamination (e.g., seed mix-ups) or cross-pollination was ruled out by examining four seeds harvested from a single plant grown in a walk-in growth cabinet under controlled, axenic conditions. These observations were further validated through repeated experiments, eliminating the likelihood of protein extraction–related artifacts.

In many cases, more than one protein biotype was detected within a single genotype. Only a small subset of genotypes exhibited biotypes in equal proportions (50:50), whereas most showed unequal frequencies. This imbalance raises concerns regarding the potential loss of one biotype over successive generations, with possible consequences for seed quality and varietal uniformity. Such genotypes represent mixtures in which allele frequencies may not be in equilibrium. Given the implications for peanut breeding and seed purity, these findings were initially discussed in 2023 with leading peanut geneticists at the University of Georgia, as well as collaborators at N.C. State University and the University of North Carolina, during meetings and scientific presentations [personal communications with Jeffrey Dunne and Ryan Andres (NCSU); David Bertioli and Soraya Leal-Bertioli (UGA); Edwin Kim and Mike Kulis (UNC), 2023.

Subsequently, similar patterns were observed in the introgression line (IL) population (F_11_) derived from *A. diogoi* and in ‘Gregory,’ where a substantial proportion of lines exhibited multiple protein biotypes (31.25% in year 1 and 19.28% in year 2) (Fig. **S4c**). However, because these genotypes were grown in the field (in 2019 and 2021) and seed protein composition is known to be environmentally sensitive, part of the observed variation may reflect environmental effects and/or cross-pollination, which can reach up to 10% (Lamon et al., 2026a). The possibility of seed mixtures, however, was ruled out because exceptional care was taken during the planting and harvesting of these lines. In contrast, the genetic heterogeneity observed for nematode resistance in ‘Tifguard’ resulted from the loss of the *Arachis cardenasii*-derived chromosome introgression on A09, likely caused by seed mixture and cross-pollination (Lamon et al., 2026a).

Protein biotype variation was also assessed in a RIL population (F_7_) derived from the cross N C 25 × ‘Georgia Green,’ evaluated in 2019 and 2021. A considerable fraction of lines displayed variable numbers of protein biotypes (30.68% in year 1 and 19.88% in year 2) (Fig **S4c**). These observations were further substantiated by analyzing multiple seeds (4–12) from individual plants selected for a desirable protein biotype. In some cases, the lower-abundance biotype was no longer detectable, whereas in others, novel biotypes emerged (Fig **S4c**). Collectively, these results suggest that homoeologous recombination–driven changes can occur rapidly (potentially detectable within a single generation) and at a higher frequency.

Seeds representing distinct protein biotypes from IL and RIL populations were collected for genotyping using low-pass sequencing–based approaches at HudsonAlpha and AgriSeq (Thermo Fisher Scientific), further supporting the presence of these biotypes. In contrast, diploid peanut accessions, such as *A. diogoi* GKP10602, consistently lacked protein biotypes, underscoring the role of polyploid genome structure in generating this variation (Lamon et al. 2025a, b).

The manifestation of biotypes as a phenotypic trait likely depends on the underlying genetic architecture of the trait. Complex traits such as seed weight and size are governed by numerous loci with small individual effects; therefore, substantial phenotypic divergence would require coordinated allele replacement or gene conversion across many genes. In contrast, traits controlled by a limited number of structural or regulatory genes—particularly those belonging to multigene families or residing in recombination hotspots—are more likely to exhibit detectable biotypic variation (see below for molecular insights).

Our findings are largely consistent with reports by Bertioli and colleagues and highlight the remarkable genetic plasticity of polyploid peanut (Lamon et al. 2025a, b). While this flexibility may confer adaptive advantages, it also poses challenges for breeding efforts aimed at developing genetically stable peanut lines, including those with reduced allergenicity. We hypothesize that such regions may eventually stabilize when homoeologous alleles converge toward either A- or B-subgenome states (Fig. **S4a**). However, this process could generate autoploid-like genomic segments within an allopolyploid background, potentially promoting uncontrolled homoeologous recombination and leading to genetic rearrangements such as translocations, deletions, insertions, and chromosome arm losses (Lamon et al. 2025a, b).

Among the 17 released varieties examined 52.9% exhibited two to five biotypes (Table **S1**). Among them, seven varieties exhibited two biotypes, with six showing a 25:75 ratio and one displaying a 50:50 ratio. No association was detected between biotype frequency and market type, year of release, or release environment. Recently, biotypes affecting traits such as grain weight, seed size, pod size, reticulation, and chlorophyll content have also been reported in released varieties (Lamon et al. 2025a). Notably, biotypes in the cultivar Tifrunner were detected at lower frequencies, potentially reflecting the greater genetic complexity underlying these traits. The presence of more than two biotypes in some cases may be attributable to aneuploidy, as reported by Lamon et al. (2025a).

Molecular insights from synthetic allopolyploid wheat and other allopolyploids, including *Brassica*, *Arabidopsis suecica*, banana, rice, and peanut, have revealed that homoeologous chromosomal exchanges (HEs) are far more prevalent than previously anticipated (Zhang et al., 2020). These events predominantly occur within gene bodies, likely due to conserved sequence similarity between homoeologues in coding regions. In contrast, homologous recombination is more often reported in promoter and terminator regions, possibly because the scarcity of polymorphisms in coding regions makes detection of these events difficult. Specific sequence motifs, such as CCN in wheat and CTT in *Brassica*, are associated with these recombination events (Zhang et al., 2020). The enrichment of HEs within coding regions, combined with the relatively high divergence between homoeologous genes (compared to homologues), has given rise to novel “recombinant” genes in wheat, *Brassica*, banana, and rice. These include chimeric transcripts that fuse the promoter of one species with the coding region of another, generating new protein variants through in-frame fusion of homoeologous genes.

In peanut, the median resolution of HE junctions spans approximately 7 kb due to an insufficient number of markers, and 65.3% of junctions are located within genic regions. The high frequency of HEs in peanut can be attributed to long stretches of DNA with high sequence similarity, resulting from the relatively recent divergence of the two progenitor species (2.2 million years ago), as well as the absence of a *Ph1*-like mechanism. Factors such as long exons and high GC content in genic regions further enhance the likelihood of recombination, promoting genetic diversity but reducing genome stability—particularly for the *Ara h1, Ara h2, Ara h3,* and *Ara h6* genes (Table **S2**). This can result in the production of restructured proteins or their complete absence, potentially due to restructuring of gene bodies, silencing of *trans*-regulatory elements and/or restructuring or rearrangement of *cis*-regulatory elements.

More recently, to facilitate the detection of homoeologous exchanges, double reduction, and tetrasomic inheritance, a synthetic tetraploid (neoallotetraploid) peanut produced by crossing *Arachis magna* with *Arachis stenosperma* was crossed with a natural tetraploid peanut (Lamon et al., 2026b). This study suggested that approximately 12% of the progeny exhibit double reductions, with such events driven by HEs primarily restricted to telomeric regions. It further confirmed that the frequency of HEs is higher than previously thought and helps explain the large number of protein biotypes and the frequent emergence of new biotypes, leading to substantial genetic diversity and trait instability (Fig **S4a-d**).

Given this complexity, maintaining the genetic purity of important germplasm may be better achieved through vegetative propagation. However, several questions remain unresolved, such as whether these variations are confined to homoeologous or tetrasomic recombination events during meiosis, or whether they can also arise during vegetative propagation or tissue culture through mitotic recombination and DNA repair involving homoeologues, thereby contributing to somaclonal variation. Comparable protein biotypes have been reported in wheat varieties and landraces (Mecham et al., 1985; Metakovsky et al., 1993) (Fig. **S4d**). However, wheat possesses an active *Ph1* gene system that suppresses homoeologous recombination (Saripalli et al., 2026). Consequently, protein biotypes in wheat are generally attributed to intentional or unintentional genotype mixtures rather than to homoeologous exchanges. However, He et al. (2017) confirmed the presence of homoeologous exchanges in wheat, and a more recent study by Zhang et al. (2020) involving synthetic wheat reconfirmed these earlier observations.

### Perspective

It is evident from studies across multiple crop species that both the timing and duration of stress strongly influence leaf, stem, and seed protein content and composition (Wang et al., 2024). Heat stress can alter the transcription of heat shock proteins in leaves within just 30 minutes of exposure. When stress occurs during seed filling, its effects on the grain proteome are more severe and lasting, often shortening the grain-filling period, reducing grain size, and increasing total protein content. In cereals—particularly wheat—high temperatures during grain filling increase overall protein accumulation but decrease metabolism-related proteins while increasing stress-defense proteins (Juhasz et al., 2018). Heat stress has also been associated with elevated levels of highly immunogenic α- and ω-gliadins and high-molecular-weight glutenins, along with reduced levels of low-molecular-weight glutenins (Juhasz et al., 2018). In contrast, cold stress during grain filling shows the opposite trend, reducing the levels of celiac-causing gliadins. Collectively, these findings suggest that heat stress tends to increase the immunogenic potential of wheat grains, whereas cold stress can reduce it.

Different crops appear to orchestrate changes in seed protein composition through distinct mechanisms, resulting in either increased or decreased allergen content. For example, in soybean, tolerant genotypes tend to maintain allergen levels, whereas susceptible genotypes may reduce them under stress. In peanut, allergen content may decrease in heat-tolerant genotypes; however, these genotypes rely on different mechanisms of heat tolerance—such as alterations in lipid composition and undisturbed chlorophyll accumulation (ICGS76) or maintenance of pollen viability (PI 493581)—which may not directly affect seed proteome composition (Selvaraj et al., 2011; Spivey et al., 2023; Ingole et al., 2025). It is also important to note that most studies to date have examined heat effects on specific proteins rather than the full seed proteome. Thus, current conclusions may reflect only pieces of a larger puzzle. More comprehensive, proteome-wide studies are needed, particularly for breeding programs that must evaluate trait stability under diverse environmental stresses and their effects on allergenicity, not just yield (Kakati et al., 2026). Additionally, crop-specific biological considerations—such as tetrasomic recombination in peanut and its influence on seed protein composition—must be accounted for.

Additionally, the impact of management practices was studied to a limited extent, given the understanding that genetic regulation of proteins is influenced by soil nitrogen content and the plant’s nitrogen status. For example, the GCN4 box, a cis-regulatory element of prolamin seed storage protein genes, senses nitrogen levels and influences seed protein content (Muller and Knudsen, 1993). Seeds derived from highly managed environments may exhibit high protein content for a few generations before these effects diminish (often described as a “legacy” or “memory” effect). Sulfur content has also been reported to have a similar influence, although the mechanism is less well understood (Rustgi et al., 2021).

As peanut allergy incidence continues to rise globally, understanding the full range of factors influencing peanut seed composition, including human interventions and environmental stressors, is increasingly urgent. Integrating genomic and proteomic tools to analyze peanut storage protein accumulation offers a foundation for developing reduced-allergen peanuts suitable for dietary therapies. With global temperatures projected to increase, predicting how elevated heat will affect peanut grain composition—from agronomic traits to immunogenic properties—becomes especially critical.

Findings from this abbreviated study suggest that certain allergenic proteins may decrease under heat stress, although direct immune assays are still needed to confirm whether heat stress reduces immunogenicity or simply degrades immunogenic protein while leaving epitopes intact. While reductions in allergenic proteins are promising, associated tradeoffs such as yield loss and potential nutritional declines temper this optimism. Nevertheless, these results highlight an opportunity to select for peanut genotypes that combine tolerance to environmental stress with reduced-allergen phenotypes observed in some susceptible varieties.

## Author contribution

**Hrishikesh Ingole & Zachary Jones:** Writing-original draft, Methodology, Investigation, Formal analysis, Data curation, Validation and Visualization. **Sachin Rustgi:** Writing-review and editing, Conceptualization, Funding acquisition, Project administration, Resources, Software, Supervision.

## Acknowledgements

The authors wish to acknowledge the receipt of seeds for this study from Dan Anco (Clemson University), Jeffrey Dunne and Ryan Andres (NCSU), Soraya Leal-Bertioli (UGA), Shyam Tallury (USDA-PGRCU), and USDA-GRIN, as well as the assistance provided by Edwin Kim, Mike Kulis, and Andrew Turner (UNC) with Western blotting.

## Data availability statement

All data presented in this study are available in the manuscript and in its supplementary information file.

