## Supplementary tables and figures for "Impacts of Anthropogenic, Environmental, and Genetic Factors on Peanut Seed Protein Content and Composition"

| **Population** | **% lines showing protein biotypes** |
| --- | --- |
| U.S. peanut mini-core | 11.5% |
| IL population (‘Gregory’ x *A. diogoi)* | 31.25% (2019), 19.28% (2021) |
| RIL population (‘Georgia Green’ x N C 25) | 30.68% (2019), 19.88% (2021) |
| Tested peanut varieties/breeding lines | 52.9% |

**Supplementary Tables**

**Table S1. Percentage of lines showing protein biotypes in various peanut populations.**

**Table S2: Comparison of genomic features across Ara h genes and the peanut genome.**

| **Genomic features** | ***Ara h* genes** | **Whole genome** |
| --- | --- | --- |
| GC content (%) | 42.90% | 36.50% |
| Average exon length (bp) | 2679.25 bp | 281 bp |
| Exon length range (bp) | 438-2205 bp | 2-1662 bp |
| Average intron length (bp) | 3249.5 bp | 618 bp |

**Supplementary Figures**

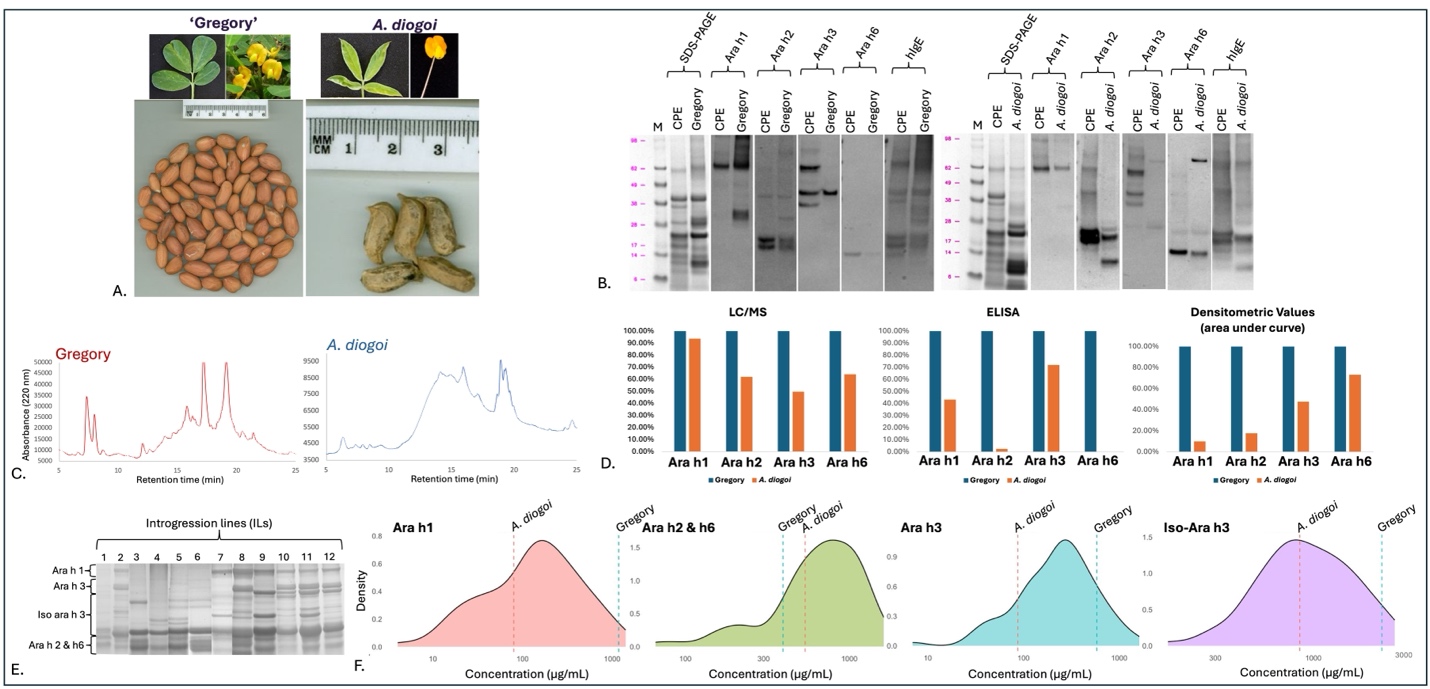

**Fig. S1. Comparison of allergenic protein content in peanut cultivar ‘Gregory’ and the wild peanut relative *Arachis diogoi* accession GKP 10602 (PI 276235).** **A.** Seed, flower, and leaf images of ‘Gregory’ and *A. diogoi*, reflecting morphological differences between the two species. **B.** SDS-PAGE and Western blot analyses for Ara h1, Ara h2, Ara h3, Ara h6, and IgE (derived from the serum of a peanut-allergic individual) in ‘Gregory’, *A. diogoi*, and peanut crude protein extract (CPE; used as a loading control/standard). **C.** Chromatograms of total seed protein extracts from ‘Gregory’ and *A. diogoi* obtained using reversed-phase high-performance liquid chromatography (experiments performed as described in Alam et al., 2026). **D.** Comparison of LC/MS, ELISA, and SDS-PAGE quantification of Ara h proteins in ‘Gregory’ and *A. diogoi* (experiments performed as described in Alam et al., 2026). E. SDS-PAGE of total seed proteins from selected introgression lines of the ‘Gregory’ × *A. diogoi* population, showing variable accumulation of major allergenic proteins. F. Frequency distribution plots showing the distribution of Ara h1, Ara h2 and Ara h6, Ara h3, and iso-Ara h3 in 87 ILs (densitometric data plotted after log transformation using R).

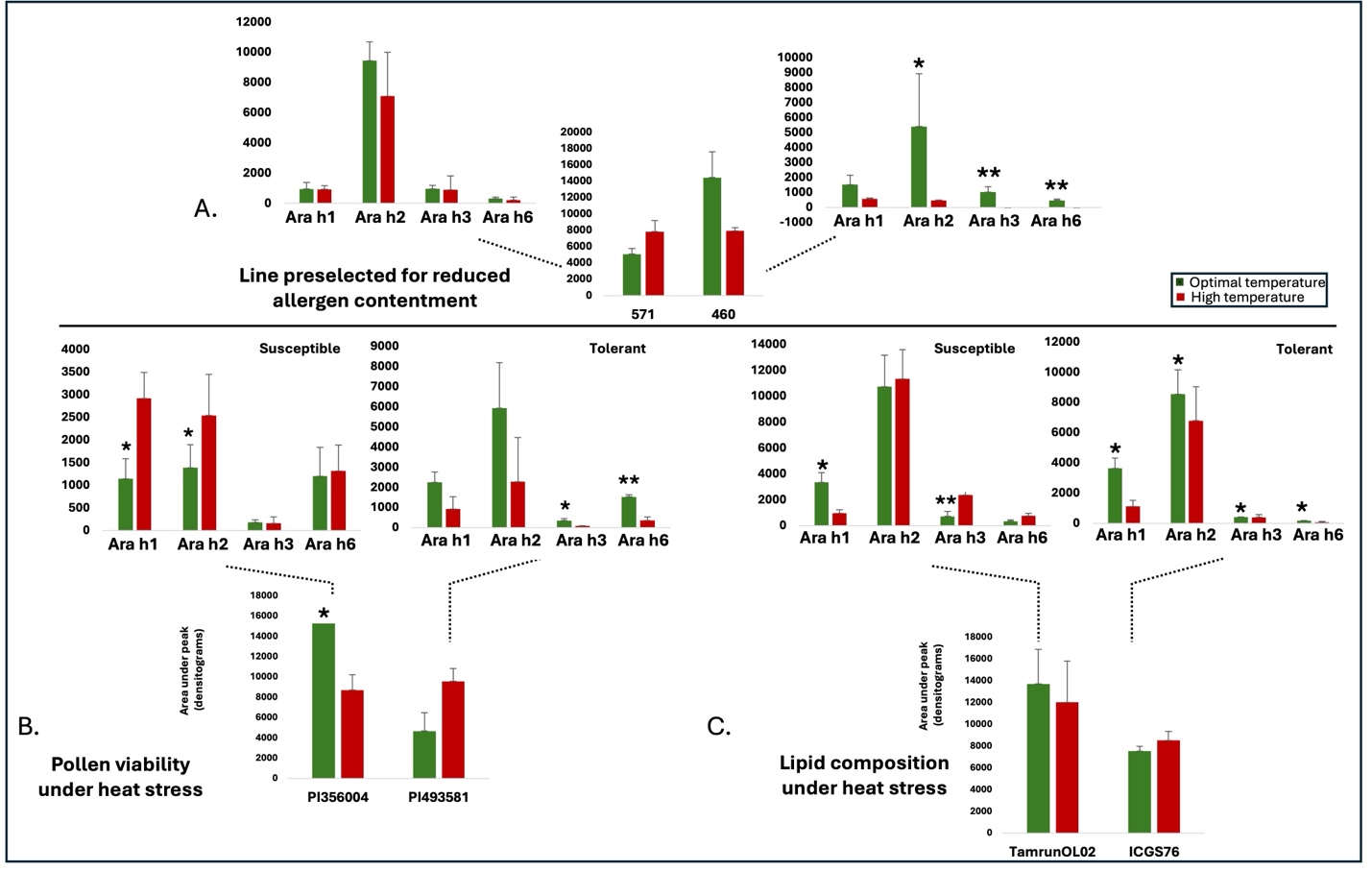

**Fig. S2. Effect of heat stress on total seed protein content and composition of major immunogenic proteins (Ara h1, Ara h2, Ara h3, and Ara h6) in known heat-tolerant and heat-susceptible peanut genotypes, as well as reduced-allergen peanut lines.** **A.** Profiles of Ara h proteins and total seed proteins in selected lines from the NC-25 × ‘Georgia Green’ peanut mapping population. **B.** Quantification of Ara h proteins and total seed proteins in lines identified as heat-susceptible or heat-tolerant based on pollen viability assays under heat stress. **C.** Quantification of Ara h proteins and total seed proteins in lines identified as heat-susceptible or heat-tolerant based on leaf lipid profiling under heat stress.

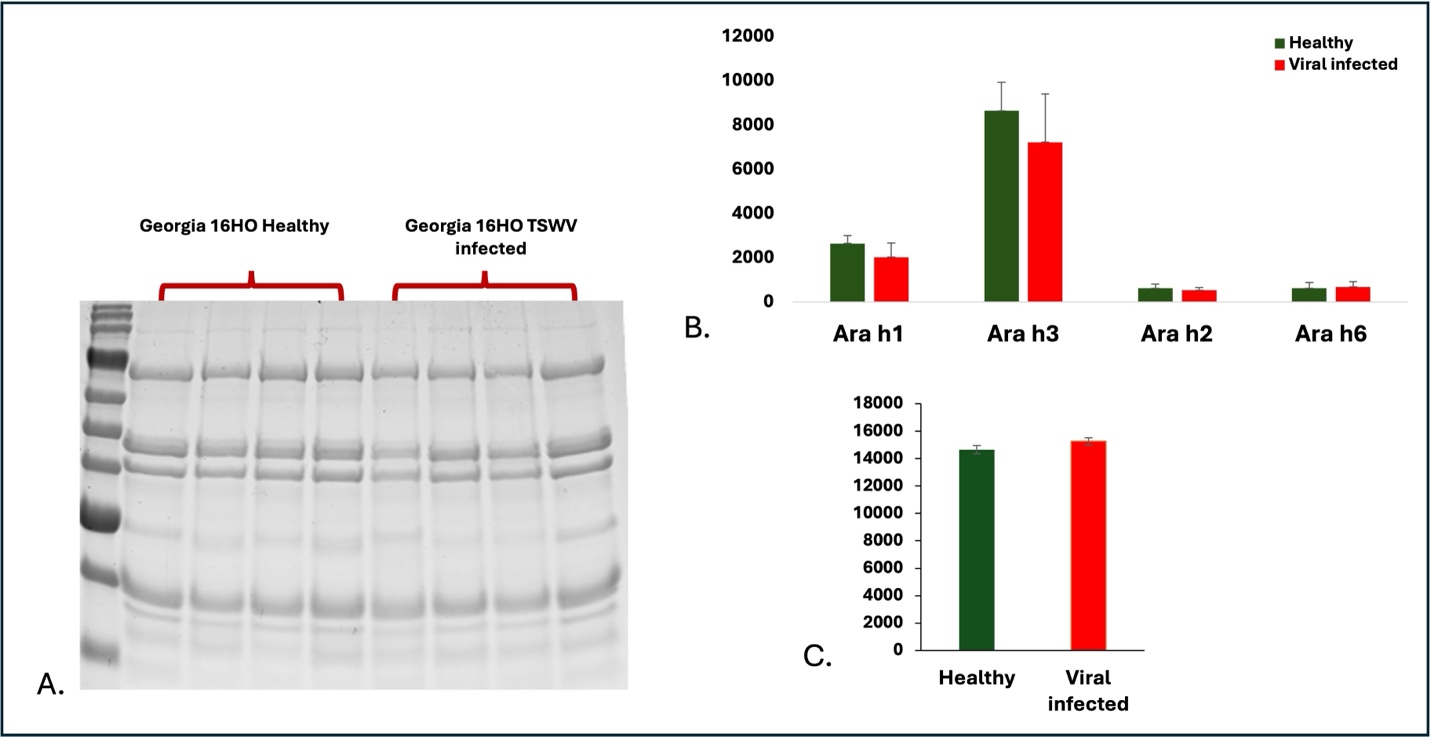

**Fig. S3. Effect of viral infection on peanut seed storage protein composition.** **A.** SDS-polyacrylamide gels of seed protein extracts from field-grown ‘Georgia 16HO’ peanut plants, comparing healthy plants and those infected with Tomato Spotted Wilt Virus (TSWV). **B.** Comparison of Ara h protein content between healthy and virus-infected plants. **C.** Total seed protein content in healthy and TSWV-infected plants.

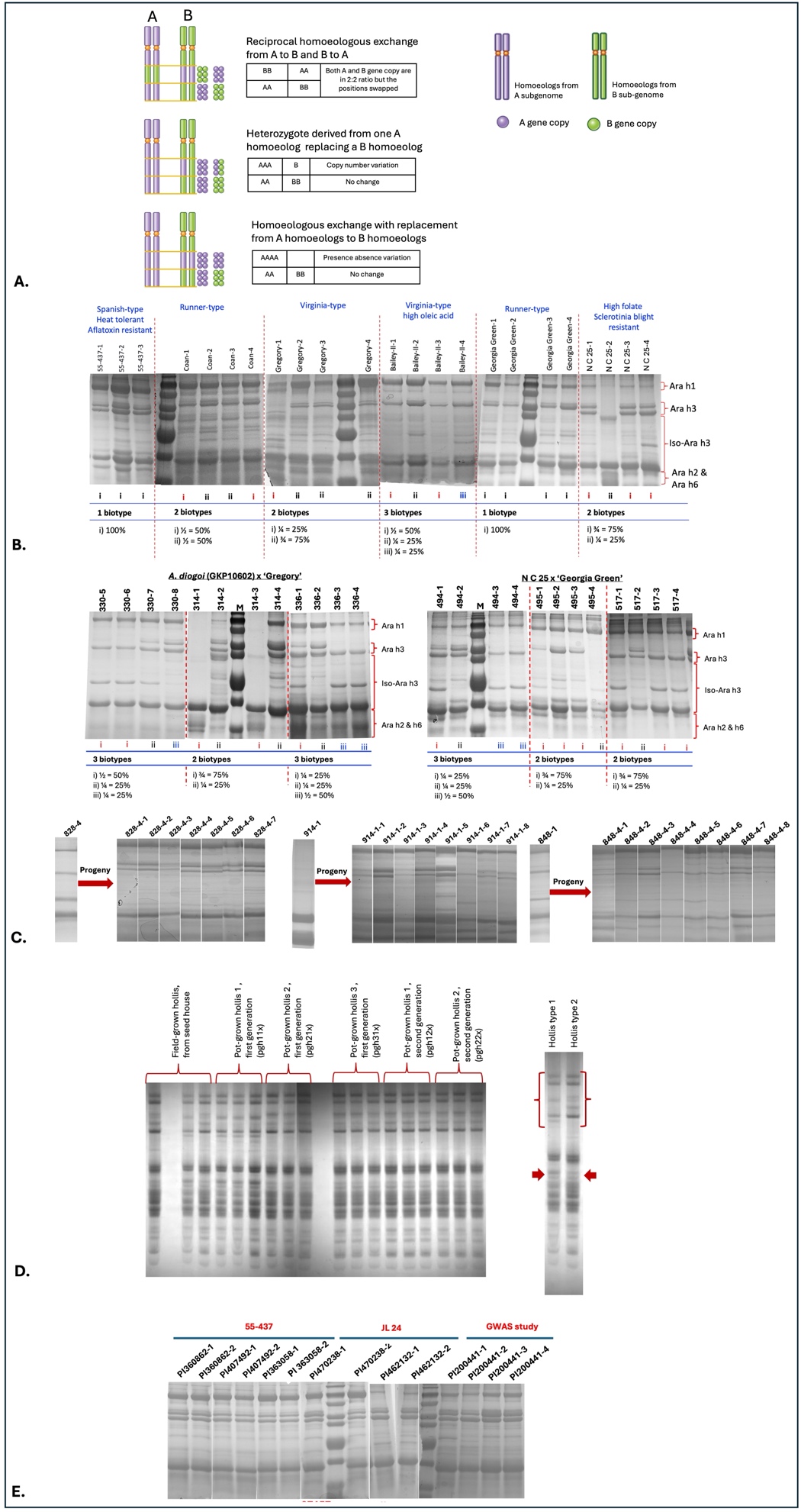

**Figure S4. Homoeologous chromosome exchange and its effect on Ara h protein profiles in peanut.** **A.** Schematic representation of homoeologous recombination in peanut and its possible outcomes (modified from Deb et al., 2023). **B.** Identification of Ara h protein biotypes using SDS-PAGE in different released peanut varieties. **C.** Identification of Ara h protein biotypes using SDS-PAGE in two peanut mapping populations (top) and their persistence in greenhouse-grown progeny (bottom). **D.** Presence of protein biotypes in wheat gliadin proteins in the released variety ‘Hollis’. **E.** Stable seed storage protein profiles in three distinct varieties, indicating a possible regulatory mechanism underlying trait stability.
